# Identifying SARS-CoV-2 entry inhibitors through drug repurposing screens of SARS-S and MERS-S pseudotyped particles

**DOI:** 10.1101/2020.07.10.197988

**Authors:** Catherine Z. Chen, Miao Xu, Manisha Pradhan, Kirill Gorshkov, Jennifer Petersen, Marco R. Straus, Wei Zhu, Paul Shinn, Hui Guo, Min Shen, Carleen Klumpp-Thomas, Samuel G. Michael, Joshua Zimmerberg, Wei Zheng, Gary R. Whittaker

**Author notes:** These authors contributed equally to this work (Catherine Z. Chen, Miao Xu). To whom correspondence should be addressed: Wei Zheng, Ph.D., Gary Whittaker, Ph.D.

## Abstract

While vaccine development will hopefully quell the global pandemic of COVID-19 caused by SARS-CoV-2, small molecule drugs that can effectively control SARS-CoV-2 infection are urgently needed. Here, inhibitors of spike (S) mediated cell entry were identified in a high throughput screen of an approved drugs library with SARS-S and MERS-S pseudotyped particle entry assays. We discovered six compounds (cepharanthine, abemaciclib, osimertinib, trimipramine, colforsin, and ingenol) to be broad spectrum inhibitors for spike-mediated entry. This work should contribute to the development of effective treatments against the initial stage of viral infection, thus reducing viral burden in COVID-19 patients.

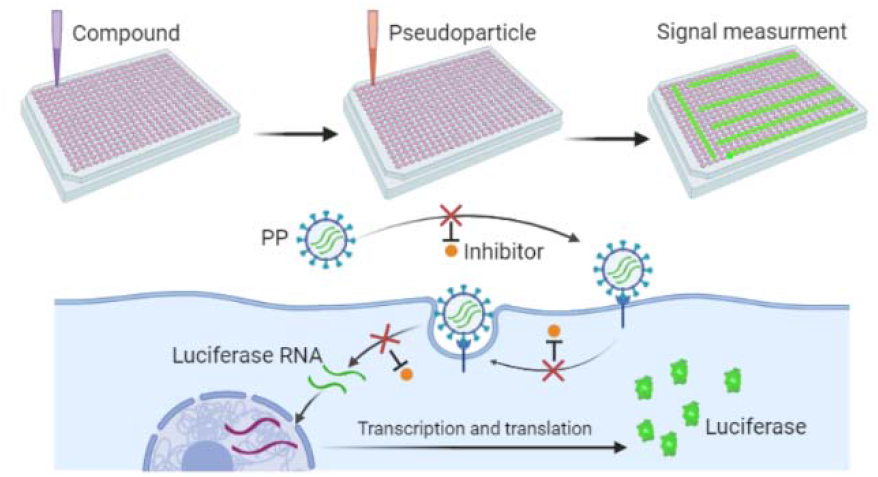

## Introduction

Coronaviruses are enveloped, single-stranded, positive-sensed RNA viruses. While some coronaviruses cause the common cold, others are highly pathogenic and have led to several outbreaks in recent years [1]. In 2003, the coronavirus strain SARS-CoV caused severe acute respiratory syndrome outbreak in Asia [2]. In 2013, the Middle East respiratory syndrome (MERS) emerged with similar clinical symptoms as SARS, and the causative agent was named MERS-CoV [3]. The <u>co</u>rona<u>vi</u>rus <u>d</u>isease 20<u>19</u> (COVID-19) was first identified in December 2019, and is caused by SARS-CoV-2, which was named based on sequence similarities to SARS-CoV [4]. While many clinical trials are actively under way for treatment of COVID-19, only remdesivir has gained emergency use authorization from the United States Food and Drug Administration. However, it is already clear that this drug alone is not enough to combat the COVID-19 pandemic [5]. Therefore, there is an unmet medical need to identify additional drugs with anti-SARS-CoV-2 activity to ameliorate disease in hundreds of millions of yet infectible individuals.

SARS-CoV-2 is a biological safety level 3 (BSL-3) pathogen. Currently, most facilities for high-throughput screening (HTS) are only BSL-2, and few BSL-3 facilities have some HTS capabilities. Several drug repurposing screens for SARS-CoV-2 have been reported, but the throughput and assay type were limited due to biocontainment requirements. Therefore, the development of BSL-2 compatible SARS-CoV-2 compound screening assays is an alternative and more facile approach for HTS and drug development. Viral entry assays utilizing pseudotyped particles (PP) are one type of cell-based BSL-2 viral assays that could be utilized for this purpose. PP contain viral envelope proteins, but carry a reporter gene instead of the viral genome, and thus display the necessary viral coat proteins for host receptor and membrane interactions without the capacity for replication. These BSL-2 viral entry assays have been successfully applied to HTS campaigns for several viruses such as Ebola virus [6], influenza [7], and human immunodeficiency virus (HIV) [8].

For SARS-CoV and MERS-CoV, the spike proteins (S) are responsible for host receptor binding and priming by host proteases to trigger membrane fusion. Thus, SARS-CoV and MERS-CoV spike proteins were pseudotyped with murine leukemia virus (MLV) gag-pol polyprotein to form SARS-S and MERS-S PP carrying luciferase reporter RNA [9,10]. The PP entry assays include inoculation of susceptible cells with SARS-S or MERS-S PP, incubation to allow luciferase reporter gene expression, and measurement of luciferase reporter activity. These protocols were successfully optimized and miniaturized in 1536-well plate formats suitable for HTS. Here, we report parallel drug repurposing screens using SARS-S and MERS-S PP entry assays to identify a set of broad-spectrum coronavirus entry inhibitors. SARS-CoV-2 live virus cytopathic effect (CPE) assay was used to test the generality of these coronavirus entry inhibitors, confirming inhibition of SARS-CoV-2 entry.

## Results

### Optimization and miniaturization of SARS-S and MERS-S PP entry assays

Both SARS-S and MERS-S PP were generated by a three plasmid co-transfection to yield particles containing capsid protein of murine leukemia virus (MLV), spike protein (SARS-S or MERS-S), and luciferase RNA (Fig. 1a). The original entry assays were developed in 24-well plates in which host cells were inoculated with PP. Upon cell entry, the particle releases the luciferase RNA reporter for subsequent expression of the luciferase enzyme (Fig. 1b) [9,10].

**Figure 1.**
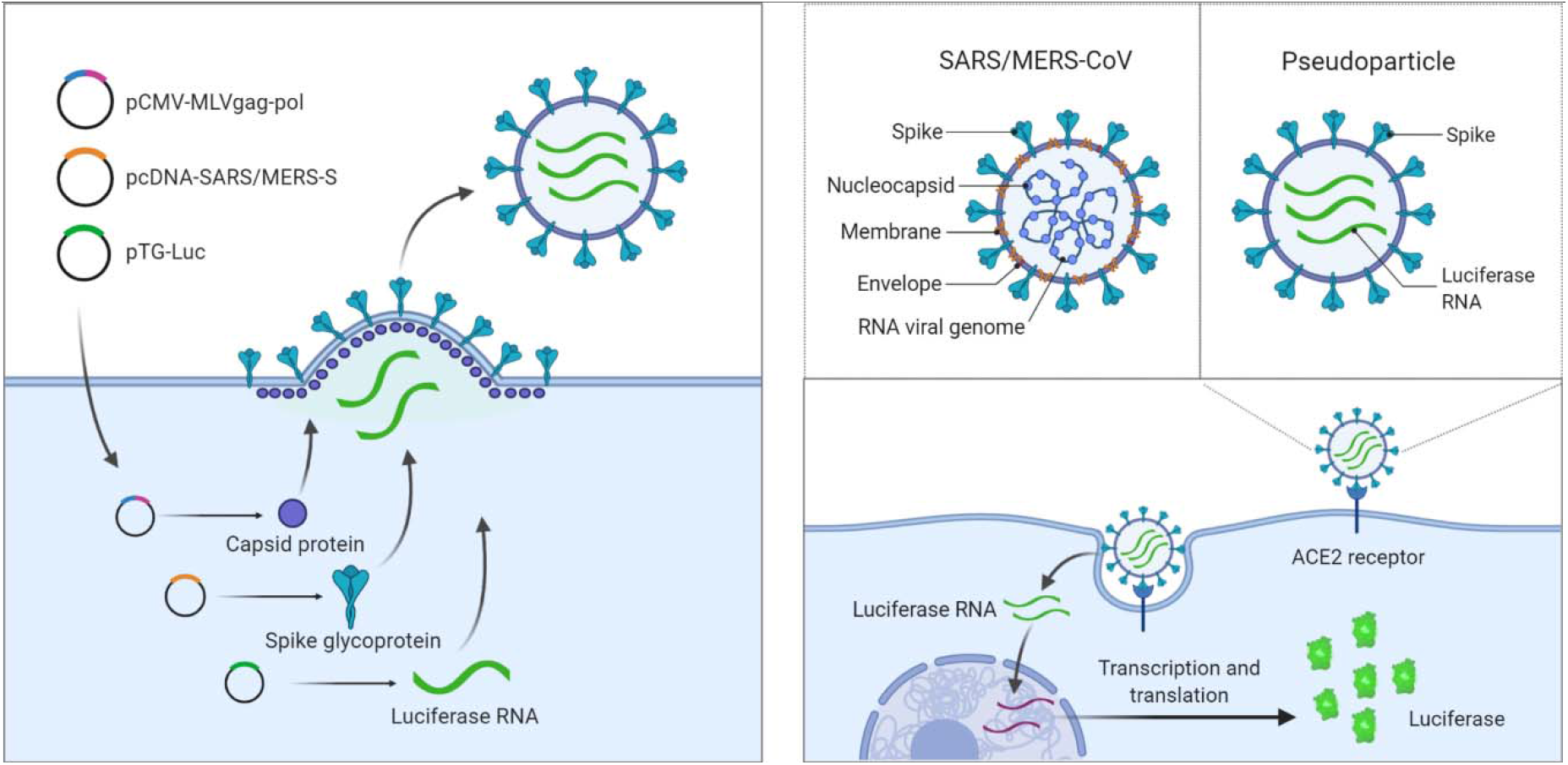
Illustration of pseudotyped particle generation and entry assay. (a) Three plasmids (pCMV-MLVgag-pol, pcDNA-SARS-S/MERS-S, and pTG-Luc) are co-transfected into HEK-293T/17 cells. The plasmids express MLV core gag-pol polyprotein, coronavirus spike glycoproteins, and luciferase RNAs, which together assemble into pseudotyped particles. (b) Comparison of SARS/MERS-CoV and pseudotyped particle, showing shared spike proteins to facilitate entry into target cell. Once cell entry occurs, RNAs of pseudotyped particles are released into cell, where they are reverse transcribed into DNAs, integrated into the genome, and express luciferase reporter enzyme. Illustrations were made with BioRender.

To optimize these assays for miniaturization into 1536-well plates, we first tested the SARS-S and MERS-S PP entry in three cell lines: Vero E6, Huh7, and Calu-3. We found that Vero E6 cells produced the highest luciferase signal for SARS-S PP assay and that Huh7 cells yielded the highest signal for MERS-S (Fig. 2a). Vesicular stomatitis virus G glycoprotein (VSV-G) is a class III fusion protein that constitutes the sole fusogenic protein, and does not require protease priming [11]. Thus, VSV-G PP was used as a positive control and produced high signals for all three cell lines (Fig. 2a). This cell tropism data agrees with previous reports [10,12]. Based on these results, the Vero E6 cell line and Huh7 cell line were chosen for SARS-S and MERS-S PP entry assays, respectively. A time course experiment showed that higher signal-to-basal (S/B) ratio was achieved with 48 h PP incubation with all glycoprotein-containing PP compared with control PP formed by two plasmid transfection lacking envelope glycoproteins that are responsible for membrane fusion (delEnv) (Fig. 2b). Therefore, the 48 h time point was used for all following experiments.

**Figure 2.**
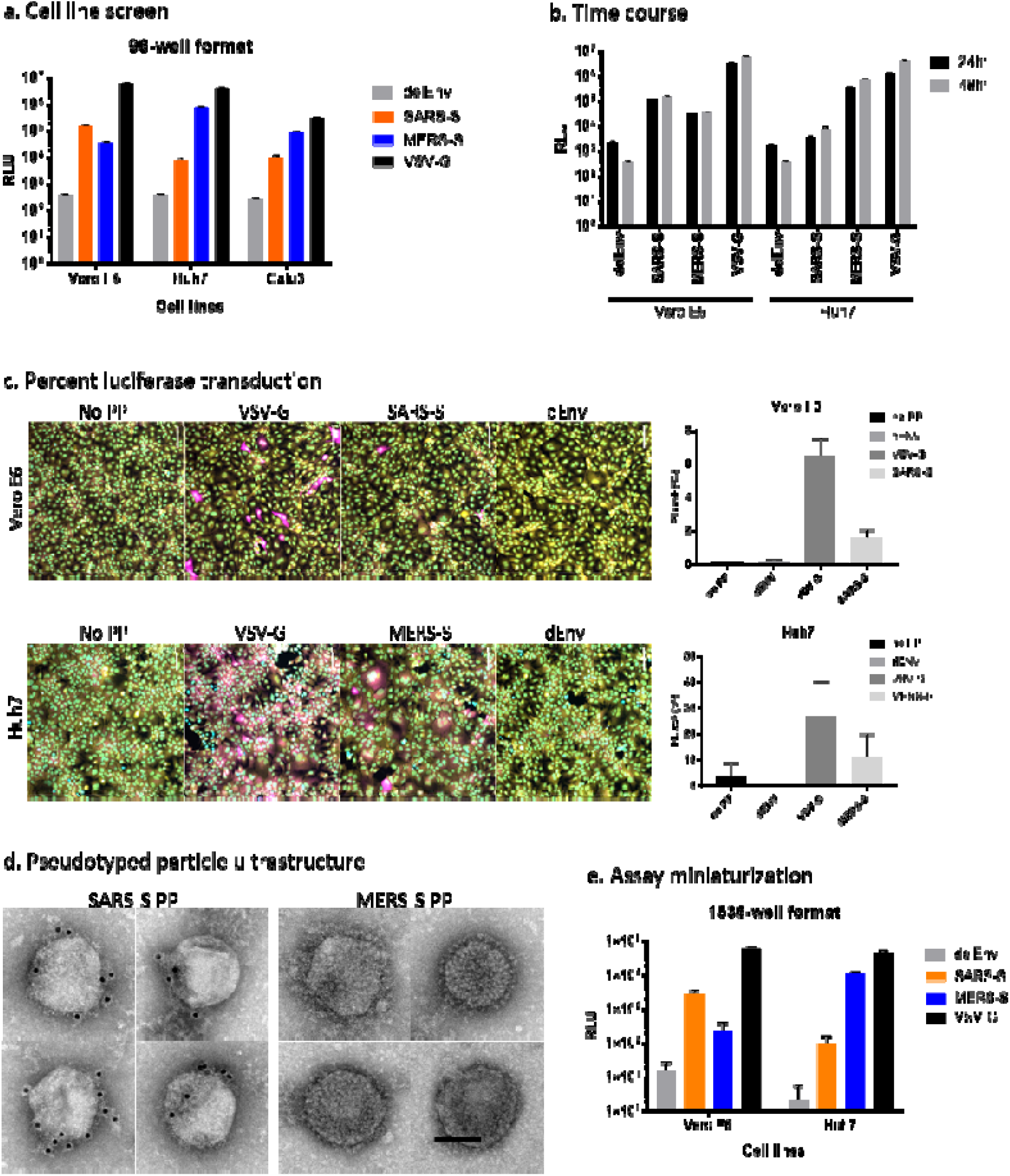
Assay optimization. (a) Entry of SARS-S, MERS-S, delEnv and VSV-G pseudotyped particles (PP) in Vero E6, Huh7 and Calu-3 cells as assayed by luciferase reporter expression. RLU = relative luminescence units. (b) Cell entry time course of PP. Luciferase reporter activity is assayed at 24 h and 48 h after PP addition. (c) Representative image montage of Vero E6 and Huh7 cells treated with VSV-G, SARS-S, or delEnv PP for 72 hours and immunostained using mouse-anti-luciferase antibody (magenta). Cells were also stained with Hoechst 33342 (cyan) for nuclei and HCS Cell Mask Green (yellow) for cell bodies. Images were captured using a 20x objective. Graphs on the right panel are high-content imaging measurements of the percentage of cells that are positive for luciferase expression. Positive cells were identified using a cell intensity threshold and the number of transfected cells was divided by the total cell count in the field. N = 9 fields per well in three wells. Error bars indicate S.D. (d) PP ultrastructure was examined by negative stain EM. Individual PP decorated with spike-like projections were observed. The presence of spike glycoproteins on the surface of SARS-S PP was confirmed by 10 nm-immunogold labeling (black dots). MERS PP displayed a dense array of spike-like projections. Scale bar = 100 nm. (e) PP entry assay was miniaturized to 1536-well format and performance of SARS-S, MERS-S, delEnv and VSV-G PP in Vero E6, Huh7 and Calu-3 cells are shown.

We examined the percentage of cells transduced with luciferase RNA by PP entry using immunofluorescence staining of luciferase protein and found that in Vero E6 cells, SARS-S PP and VSV-G PP produced1.6% and 6.5% luciferase positive cells, respectively (Fig. 2c). In Huh7 cells, MERS-S PP and VSV-G pp transduction produced 10.9% and 26.8% luciferase positive cells, respectively (Fig. 2c). In all cells, the negative control delEnv PP and no PP conditions produced negligible luciferase staining. The percentage of luciferase positive cells correlated with luciferase enzyme activity when comparing different PP in the same cell line. However, cell line comparisons did not show correlation between each PP, which may in part reflect differences in the amount of luciferase expression per cell. The ultrastructure of SARS-S and MERS-S PP were examined by negative stain electron microscopy (EM) to ensure that they showed the expected morphology. EM analysis revealed regularly sized, 125-200 nm diameter spherical structures, that were often partially or completely covered with a dense array of fine filamentous or lollipop shaped projections, consistent with expected appearance of spike glycoproteins (Fig. 2d). The presence of SARS-S on the surface of SARS-S PP was further confirmed by immunogold labeling (Fig. 2d). MERS-S PP displayed a conspicuous dense coat of spike-like structures, but lack of a primary antibody has thus far precluded confirmation of their identity with immunogold labeling.

Both SARS-S and MERS-S PP entry assays were then miniaturized into 1536-well plate format. The cell tropism pattern in the 1536-well format matched what was seen in the 96-well format (Fig. 2e). For SARS-S PP, the best assay performance was seen in Vero E6 cells compared with delEnv PP, with S/B of 182.3, coefficient of variation (CV) of 24.1%, and a Z’ factor of 0.26. For MERS-S PP the best assay performance was seen in Huh7 cells, with an S/B of 5325.8, CV of 10.9, and Z’ factor of 0.67. Therefore, the SARS-S PP entry assay in Vero E6 cells, and MERS-S PP entry assay in Huh7 cells, were robust and advanced to HTS.

### SARS-S and MERS-S entry inhibitor drug repurposing screens

The NCATS pharmaceutical collection (NPC) of 2,678 compounds, representing approved or investigational drugs [13], was used for drug repurposing screens of both SARS-S and MERS-S PP entry assays. The primary screens were carried out at four compound concentrations (0.46, 2.3, 11.5, and 57.5 μM). Compound cytotoxicity as determined by an ATP content assay was counter screened in both Vero E6 and Huh7 cell lines, at the same concentrations (Fig. 3). All primary screening datasets were deposited to PubChem (Table 1).

**Figure 3.**
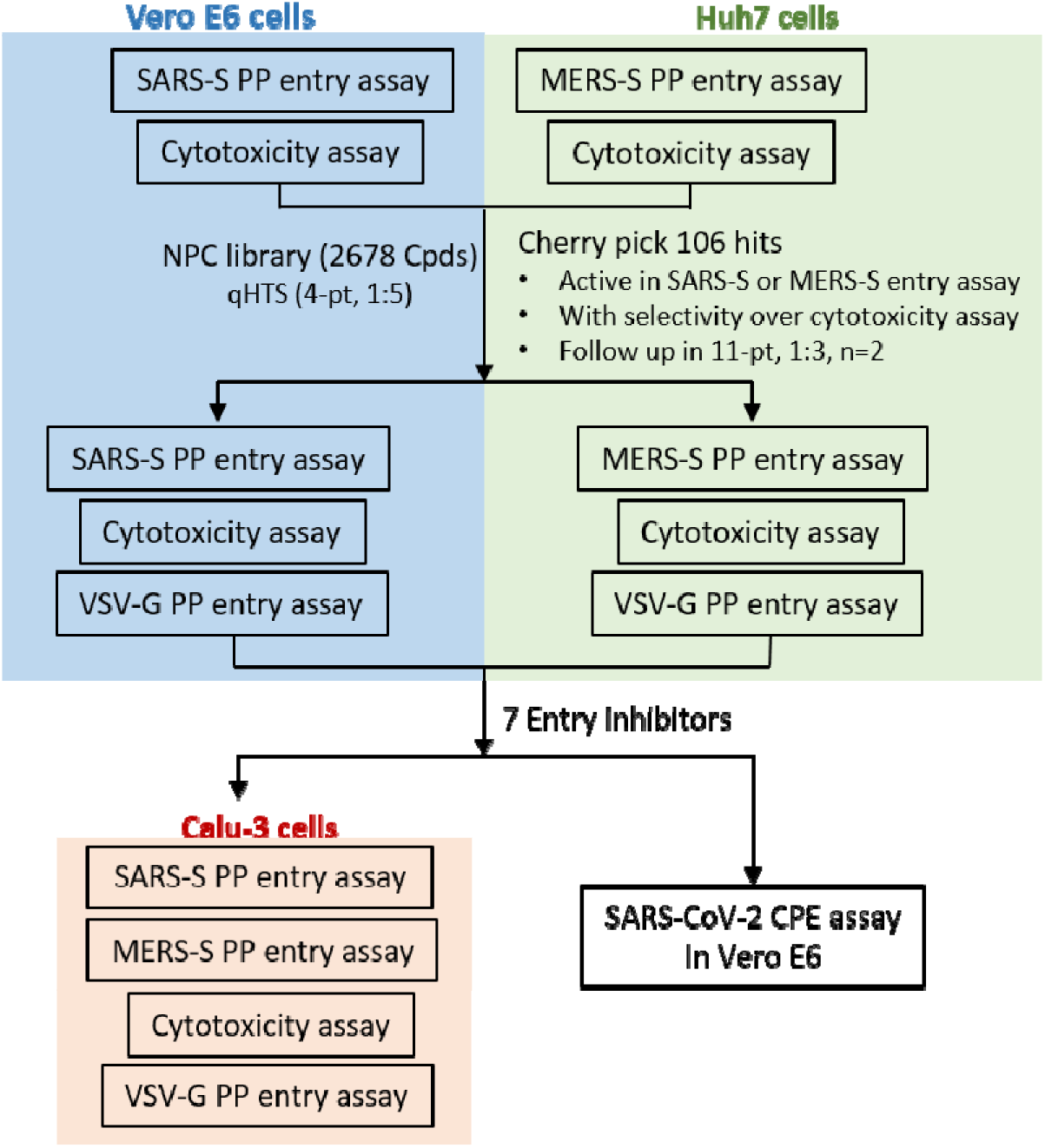
Schematic of repurposing screen and follow up assays.

**Table 1.** PubChem assay IDs (AIDs). Datasets can be found at https://pubchem.ncbi.nlm.nih.gov/ under the following AIDs.

| AID | # of Compounds | Concentration response format | Assay | Cell line |
| --- | --- | --- | --- | --- |
| 1479145 | 2678 | 4-pt, 1:5 | SARS-S PP entry | Vero E6 |
| 1479150 | 2678 | 4-pt, 1:5 | Cytotoxicity | Vero E6 |
| 1479149 | 2678 | 4-pt, 1:5 | MERS-S PP entry | Huh7 |
| 1479147 | 2678 | 4-pt, 1:5 | Cytotoxicity | Huh7 |
| 1479148 | 106 | 11-pt, 1:3 | SARS-S PP entry | Vero E6 |
| 1494158 | 106 | 11-pt, 1:3 | VSV-G PP entry | Vero E6 |
| 1479144 | 106 | 11-pt, 1:3 | Cytotoxicity | Vero E6 |
| 1494157 | 106 | 11-pt, 1:3 | MERS-S PP entry | Huh7 |
| 1494156 | 106 | 11-pt, 1:3 | VSV-G PP entry | Huh7 |
| 1479146 | 106 | 11-pt, 1:3 | Cytotoxicity | Huh7 |

The criteria to select hits for follow-up experiments include compounds in curve classes 1 and 2 with efficacy > 50% in the PP entry assay, and little to no cell killing effect in the cytotoxicity assays using Vero E6 or Huh7 cells. Sixty-one and sixty-five compounds were identified as hits from SARS-S and MERS-S PP viral entry assays, respectively. After removing 20 overlapping hits, a total of 106 primary hits (4.0% hit rate) were selected for activity confirmation and follow-up studies.

### Hit confirmation and follow up assays

In our secondary assays, we retested the 106 cherry-picked hits in the original SARS-S and MERS-S PP entry assays, along with ATP content cytotoxicity assays at 11 concentrations with 1:3 titration. PP entry assays rely on luciferase RNA reporter expression, a process which involves the reverse transcription of luciferase RNA, integration into host genome, and expression. Indeed, some of the confirmed hits had known mechanisms of action against reverse transcriptase (adefovir and tenofovir disoproxil fumarate), and integrase (elvitegravir and dolutegravir). Therefore, another counter assay, VSV-G PP entry assay, was tested in both Vero E6 and Huh7 cell lines against the 106 hits. In addition to eliminating false positives that inhibit luciferase expression, this assay identified compounds that specifically blocked spike glycoprotein-mediated PP entry. All datasets for secondary assays are publicly available on PubChem (Table 1).

These follow up assays yielded a set of 7 inhibitors that showed greater than 10-fold selectivity to either SARS-S or MERS-S PP entry assays compared with the VSV-G PP entry assays, and a safety index greater than 10 fold (CC_50_/EC_50_) (Fig. 4a, b, Table 2). Of these 7 compounds, only cepharanthine was active against both SARS-S and MERS-S with greater than 10-fold selectivity. While trimipramine, copansilib, abemaciclib and osimertinib showed some level of selectivity towards either SARS-S or MERS-S entry versus VSV-G entry, they only reached 10-fold selectivity in one of the spike PP entry assays. Ingenol and NKH477 were only active in SARS-S PP entry in Vero E6, and not in MERS-S entry in Huh7 cells.

**Figure 4.**
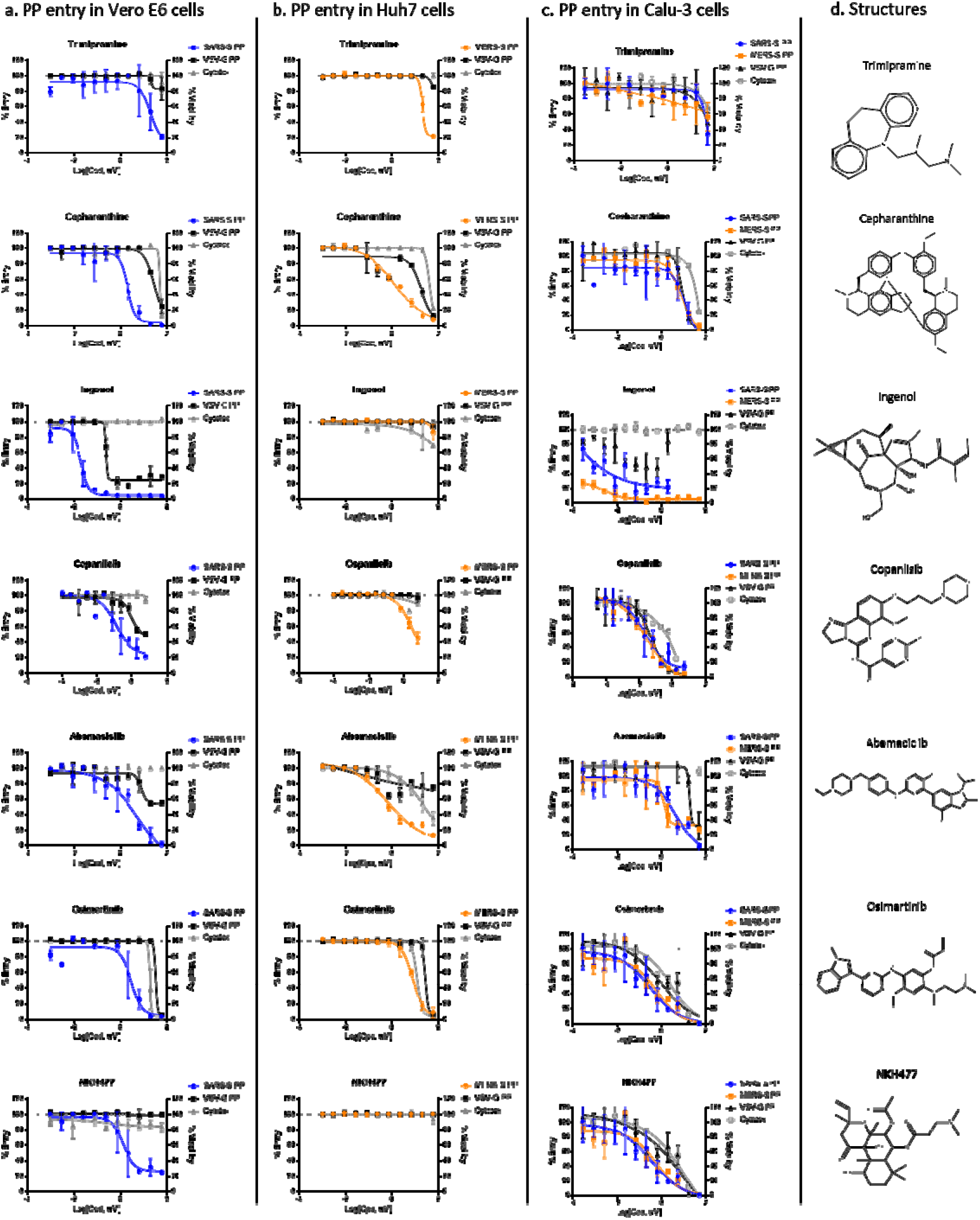
Concentration response of entry inhibitors. (a) Concentration response of entry inhibitors against SARS-S and VSV-G PP entry in Vero E6 cells. (b) Concentration response of entry inhibitors against MERS-S and VSV-G PP entry in Huh7 cells. (c) Concentration response of entry inhibitors against SARS-S, MERS-S and VSV-G PP entry in Calu-3 cells. (d) Compound structures.

**Table 2.** SARS-S and MERS-S selective compounds and their anti-SARS-CoV-2 activity. N/A = Not active, highest concentration tested is listed. MOA = Mechanism of action

| **Compound Name**<br>**(MOA)** | **SARS-S PP**<br>**in Vero E6** |  | **VSV-G PP**<br>**in Vero E6** |  | **Vero E6**<br>**Cytotoxicity** |  | **SARS-CoV-2**<br>**CPE** |  | **SARS-CoV-2**<br>**Cytotoxicity** |  |
| --- | --- | --- | --- | --- | --- | --- | --- | --- | --- | --- |
|  | **EC50**<br>**(μM)** | **Efficacy**<br>**(%)** | **EC50**<br>**(μM)** | **Efficacy**<br>**(%)** | **CC50**<br>**(μM)** | **Efficacy**<br>**(%)** | **EC50**<br>**(μM)** | **Efficacy**<br>**(%)** | **CC50**<br>**(μM)** | **Cytotox**<br>**(%)** |
| <u>NKH477</u><br>(Adenylyl cyclase activator) | 1.36 | 71.4 | N/A,<br>>57.5 | 0.0 | N/A,<br>>57.5 | 0.0 | 23.06 | 45.6 | 25.20 | 42.0 |
| <u>Trimipramine</u><br>(Tricyclic antidepressant) | 4.29 | 90.9 | 30.40 | 29.2 | 54.06 | 16.6 | 20.52 | 48.1 | >20 | 16.6 |
| <u>Osimertinib</u><br>(EGFR inhibitor) | 2.71 | 117.5 | 42.94 | 118.4 | 17.10 | 99.6 | 3.98 | 60.0 | 10.00 | 99.7 |
| <u>Ingenol</u><br>(Topical anti-tumor medication) | 0.02 | 93.3 | 0.24 | 76.4 | 0.00 | -4.0 | 0.06 | 38.2 | N/A,<br>>20 | 0.0 |
| <u>Cepharanthine</u><br>(Anti-inflammatory, antineoplastic) | 1.92 | 90.9 | 21.52 | 76.9 | 42.94 | 106.2 | 1.41 | 92.5 | 11.22 | 99.0 |
| **Compound Name**<br>**(MOA)** | **MERS-S PP**<br>**in Huh7** |  | **VSV-G PP**<br>**in Huh7** |  | **Huh7**<br>**Cytotoxicity** |  |  |  |  |  |
|  | **EC50**<br>**(μM)** | **Efficacy**<br>**(%)** | **EC50**<br>**(μM)** | **Efficacy**<br>**(%)** | **CC50**<br>**(μM)** | **Efficacy**<br>**(%)** |  |  |  |  |
| <u>Abemaciclib</u><br>(CDK inhibitor) | 0.38 | 82.3 | 0.27 | 27.1 | 17.10 | 90.0 | 3.16 | 68.7 | 7.08 | 34.0 |
| <u>Copansilib</u><br>(PI3K inhibitor) | 3.12 | 65.8 | 9.87 | 6.2 | 1.75 | 13.6 | N/A,<br>>20 | 0.0 | 5.74 | 42.4 |
| <u>Cepharanthine</u><br>(Anti-inflammatory,<br>antineoplastic) | 1.71 | 108.4 | 24.15 | 115.3 | 38.27 | 88.9 | 1.41 | 92.5 | 11.22 | 99.0 |

These 7 confirmed entry inhibitors were then tested in SARS-S, MERS-S and VSV-G PP entry assays in Calu-3 cells (Fig. 4c). While most entry inhibitors failed to show selectivity towards spike-mediated entry in Calu-3 cells, abemaciclib did show >10-fold selectivity towards both SARS-S and MERS-S based entry compared with VSV-G.

### SARS-CoV-2 cytopathic effect assay to identify broad acting coronavirus entry inhibitors

To test whether the confirmed SARS-S and MERS-S mediated PP entry inhibitors are active against SARS-CoV-2, we further tested the top 7 compounds in a SARS-CoV-2 cytopathic effect (CPE) assay [14]. We found that 6 out of 7 entry inhibitors significantly reduced (>30%) CPE caused by the SARS-CoV-2 infection in Vero E6 cells (Fig. 5, Table 2). Cepharathine was found to be active against SARS-S in Vero E6 and MERS-S in Huh7 cells, and inhibited SARS-CoV-2 CPE to near full efficacy with bell-shaped concentration response due to cytotoxicity (Fig. 5b). Five other compounds, trimipramine, ingenol, abemaciclib, osimertinib and NKH447 also protected against SARS-CoV-2 induced CPE, but to lesser degrees than cepharanthine (Fig. 5).

**Figure 5.**
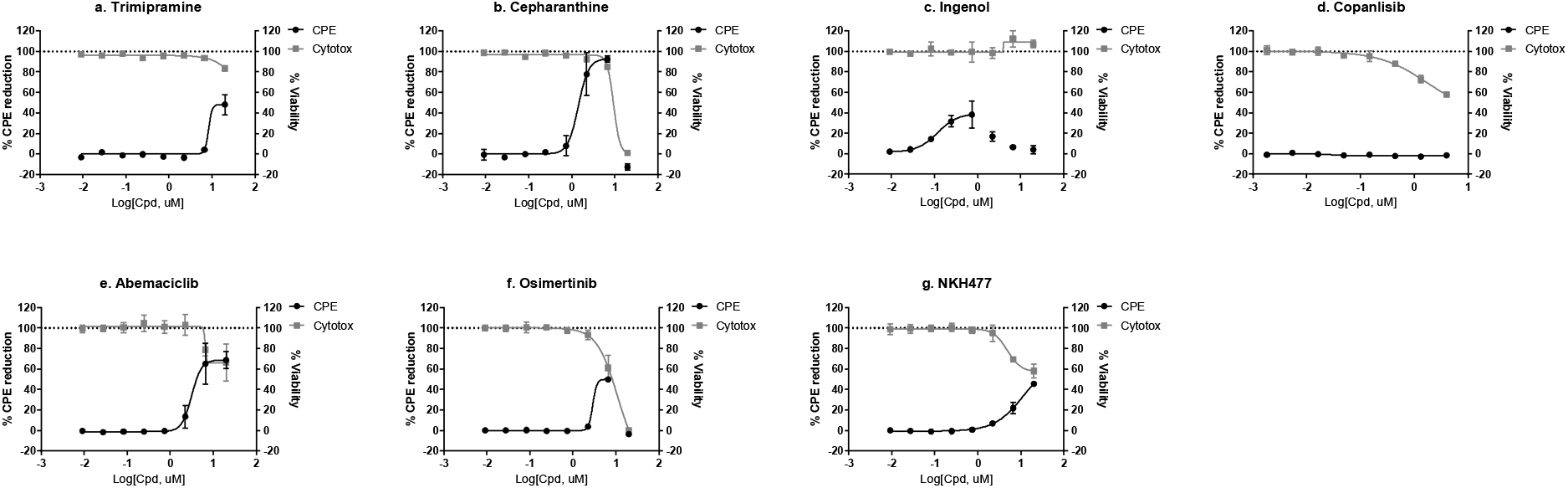
SARS-CoV-2 CPE assay and cytotoxicity concentration response for (a) Trimipramine, (b) Cepharanthine, (c) Ingenol, (d) Copanisib, (e) Abemaciclib, (f) Osimertinib, and (g) NKH477.

## Discussion

Viruses rely on host cells for replication, and cell entry is the first step of the viral infection life cycle, and a prime target for drug intervention. Both broad-spectrum and pathogen-specific inhibitors of viral entry have been proposed for emerging viruses such as Ebola virus and coronaviruses [15,16]. Proven therapeutics for viral entry include several approved drugs targeting CCR5, the host co-receptor for HIV [17]. In SARS-CoV and SARS-CoV-2, angiotensin-converting enzyme 2 (ACE2) has been recognized as a high affinity binding receptor for the viral spike glycoprotein, while dipeptidyl peptidase-4 (DPP4) is the receptor for MERS-CoV [18,19]. Following receptor binding, membrane fusion is mediated by spike protein cleavage by host cell proteases. TMPRSS2 protease has been shown to be the predominant protease in Calu-3 cells, which mediates ACE2-dependent direct membrane fusion that does not involve the endocytic pathway [12]. Alternate entry pathways are used in cell lines such as Vero E6 and Huh7 that involve endocytosis of viral particles and cathepsin protease priming for membrane fusion [18]. Here, we have applied phenotypic SARS-S and MERS-S PP entry assays for drug repurposing screens with the potential of identifying viral entry inhibitors with different mechanisms of action.

In this study, we identified 7 coronavirus spike-mediated entry inhibitors out of a library of 2,678 approved drugs (Fig. 4). After further testing in a SARS-CoV-2 live virus CPE assay and removing cytotoxic compounds, we found that 6 out of 7 entry inhibitors were able to rescue the CPE of SARS-CoV-2 infection (Fig. 5), indicating the utility of these PP entry assays. Although the exact mechanism for entry inhibition is unclear, these six compounds, inhibited SARS-S and MERS-S PP cell entry with greater potency than VSV-G PP cell entry (Fig. 4), indicating their coronavirus-specific inhibitory activities on viral entry into host cells. Of these six, only cepharanthine and abemaciclib have been reported to have anti-SARS-CoV-2 effects, while the other compounds are novel [20]. We found that cepharanthine is an inhibitor of spike-mediated cell entry and rescued the CPE of SARS-CoV-2 to full efficacy. This was corroborated by a recent study, which found that cepharanthine was able to block SARS-CoV-2 induced CPE in Vero E6 cells, only when added during the viral entry time period, and not the post-entry period [21]. Cepharanthine is a natural product used in Japan since the 1950s for treatments of several diseases without major side effects [22]. It has polypharmacology, with anti-inflammatory activity linked to AMPK activation and NFκB inhibition [22]. Cepharanthine has previously reported antiviral activities against HIV, SARS-CoV, HCoV-OC43, human T-lymphotropic virus (HTLV) and hepatitis B virus (HBV) [23].

We also identified two other approved drugs, abemaciclib and osimertinib as entry inhibitors that showed greater than 50% rescue of the SARS-CoV-2 CPE (Fig.s 4 and 5).

Interestingly, abemaciclib also showed spike-specific PP entry inhibition in Calu3 cells over VSV-G mediated PP entry (Fig. 4c). Abemaciclib is a CDK4/6 inhibitor that is approved by the FDA for breast cancer treatment [24,25]. Cyclin-dependent kinases (CDK) are a group of serine-threonine kinases that regulate the cell cycle, and have been targeted for anticancer drug development. Additionally, antiviral activities of CDK inhibitors have been reported against HIV, herpes simplex virus (HSV), HBV and Zika virus [26]. The antiviral mechanism of action for CDK inhibitors works mainly through the suppression of viral genome replication in host cells [26]. Our data suggests that abemaciclib inhibits CPE of coronaviruses by blocking cell entry in Vero E6, Huh7 and Calu-3 cells. Therefore, the structure of this compound may have the potential to be optimized as a more potent SARS-CoV-2 entry inhibitor. Osimertinib is an inhibitor of T790M mutant of epidermal growth factor receptor (EGFR), and is approved by the FDA for treatment of patients with metastatic mutation-positive non-small cell lung cancer [27]. Osimertinib does not have previous reported antiviral activities. We found it to rescue the SARS-CoV-2 CPE to 60% efficacy, albeit with a narrow therapeutic window due to cytotoxicity (Fig. 5f).

Three other inhibitors of spike-mediated PP entry were found to rescue SARS-CoV-2 CPE to less than 50% efficacy: trimipramine, ingenol, and NKH477 (Fig. 5a, c, g). Trimipramine is an oral tricyclic antidepressant. Chemically, trimipramine is a basic amine compound belonging to cationic amphiphilic drugs. The antiviral activity of trimipramine has been reported to block the viral entry for Ebola virus and influenza [28,29]. Due to its chemical property, trimipramine as a basic amine can accumulate in acidic organelles such as the late endosomes and lysosomes in cells. High concentration of basic amine drugs in late endosomes and lysosomes may block viral genome release into cytosol [28,29]. However, for coronaviruses, this effect might be more prominent in cell lines such as Vero E6 and Huh7, but not in Calu-3 cells, which has endocytosis independent entry [18]. In accordance with this, trimipramine’s entry inhibition activity was not confirmed in Calu-3 cells (Fig. 4). Importantly, the antiviral entry activity of trimipramine has not yet been reported. In addition, clomipramine, a close analog of trimipramine, was also reported to protect against SARS-CoV-2 CPE through inhibition of autophagy [14]. In the current study, clomipramine was found to be active against SARS-S PP entry and non-cytotoxic in the primary screen, but was not selected for further follow-up because its potency was below the threshold criteria. Ingenol mebutate is a cell death inducer approved by the FDA for topical treatment of actinic keratosis [30]. Due to its topical delivery route and mechanisms of action, ingenol is unlikely to be useful for treatment of COVID-19. NKH477, also called colforsin, is a derivative of forskolin and a potent activator of adenylate cyclase [31]. It is approved in Japan for multiple indications and does not have reports of direct antiviral activities.

A number of drug repurposing and computer-aided virtual screens have been reported for SARS-CoV-2. It is a common phenomenon that the potencies identified in drug repurposing are not high enough to be clinically relevant when compared to the human plasma concentrations achievable at approved dosing regimens [32]. Drug combination therapy has been proposed as a practical and useful approach for drug repurposing to treat emerging infectious diseases, as drug synergy may reduce the individual drug concentrations in the combinations. The synergistic effect of two- or three-drug combination therapy can increase the therapeutic effect, reduce the doses of individual drugs, and thus reduce potential adverse effects [32]. Ohashi H et al. has reported that the combination of cepharanthine (entry inhibitor) and nelfinavir (HIV protease inhibitor) enhanced the anti-SARS-CoV-2 activity [21]. We believe that these coronavirus specific viral entry inhibitors may have utility in a drug combination therapy with other anti-SARS-CoV-2 drugs that have different mechanisms of action, such as remdesivir (the viral RNA dependent RNA polymerase inhibitor), or lysomotropic autophagy inhibitors. In addition, considering that there have been three different coronavirus outbreaks in the past 20 years, these broad acting inhibitors of spike-mediated cell entry might also have utility in drug development efforts for possible future outbreaks.

## Methods

### Reagents

The following items were purchased from ThermoFisher: Dulbecco's Modified Eagle's Medium (DMEM) (11965092), Pen/Strep (15140), TrypLE (12604013), PBS -/- (w/o Ca2+ or Mg2+) (10010049), HCS Cell Mask Green (H32714), goat anti-mouse AlexaFluor 647 (A28175), and Hoechst 33342 (H3570). EMEM (30-2003) was purchased from ATCC. Hyclone FBS (SH30071.03) was purchased from GE Healthcare. Pseudotyped particles (PP) for SARS-S PP, MERS-S PP, VSV-G PP and delEnv PP (PP without fusion proteins) were produced by Codex Biosolutions (Gaithersburg, MD) using previously reported methods [9,10]. Microplates were purchased from Greiner Bio-One: white tissue-culture treated 96-well plates (655090), black μclear 96-well plates (655083), white tissue-culture treated 384-well plates (781073), and white tissue-culture treated 1536-well plates (789173-F). The following were purchased from Promega: BrightGlo Luciferase Assay System (E2620), CellTiter-Glo Luminescent Cell Viability Assay (G7573). ATPLite Luminescence Assay kit was purchased from PerkinElmer (6016949). Cell Staining Buffer (420201) was purchased from BioLegend. Paraformaldehyde (PFA) was purchased from Electron Microscopy Sciences (15714-S). Mouse-anti-firefly luciferase antibody was purchased from Santa Cruz (sc-74548). SARS-S antibody was purchased from BEI Resources (NR-617).

### Cell lines and cell culture

Vero E6 cells (ATCC #CRL-1586) were cultured in EMEM with 10% FBS. Huh7 cells (JCRB cell bank #JCRB0403) were cultured in DMEM with 10% FBS. Calu-3 cells (ATCC #HTB-55) were cultured in EMEM with 10% FBS.

### Pseudotyped particle (PP) entry assays

96-well format: Cells were seeded in 50 μL/well media (20,000 cells/well for Vero E6 and Huh7, and 40,000 cells/well for Calu-3 cells), and incubated at 37◻°C, 5% CO_2_ overnight (~16 h). Supernatant was removed, and 50 μL/well of PP was added. Plates were spin-inoculated at 1500 rpm (453 xg) for 45 min, incubated for 2 h at 37◻°C, 5% CO_2_, then 50 μL/well of growth media was added. The plates were incubated for 48 h at 37◻°C, 5% CO_2_. The supernatant was removed, 100 μL/well of Bright-Glo (Promega) was added, incubated for 5 min at room temperature, and luminescence signal was measured using a PHERAStar plate reader (BMG Labtech).

384-well format: 10,000 cells/well of Calu-3 cells were seeded in 10 μL media, and incubated at 37□°C, 5% CO_2_ overnight (~16 h). Supernatant was removed, 10 μL/well of 2x compounds in media was added, incubated for 1 h, before 10 μL/well PP was added. Plates were spin-inoculated at 1500 rpm (453 xg) for 45 min, and incubated for 48 h at 37◻°C, 5% CO_2_. The supernatant was removed, 20 μL/well of Bright-Glo (Promega) was added, incubated for 5 min at room temperature, and luminescence signal was measured using a PHERAStar plate reader (BMG Labtech).

1536-well format: Cells were seeded at 2000 cells/well in 2 μL media, and incubated at 37□°C, 5% CO_2_ overnight (~16 h). Compounds were titrated in DMSO, and 23 nL/well was dispensed via an automated pintool workstation (Wako Automation). Plates were incubated for 1◻h at 37C, 5% CO2, and 2 μL/well of PP was dispensed. Plates were spinoculated by centrifugation at 1500 rpm (453 xg) for 45 min, and incubated for 48 h at 37◻°C, 5% CO_2_. After the incubation, the supernatant was removed with gentle centrifugation using a Blue Washer (BlueCat Bio). Then, 4 μL/well of Bright-Glo (Promega) was dispensed, incubated for 5 min at room temperature, and luminescence signal was measured using a ViewLux plate reader (PerkinElmer). All data was normalized with wells containing PP as 100%, and delEnv PP as 0% entry.

### ATP content cytotoxicity assay

Cells were seeded at 1000 cells/well in 2 μL/well media in 1536-well plates, and incubated at 37□°C, 5% CO2 overnight (~16 h). Compounds were titrated in DMSO, 23 nL/well was dispensed via an automated pintool workstation (Wako Automation). Plates were incubated for 1◻h at 37C, 5% CO2, before 2 μL/well of media was added. Plates were incubated for 48◻h at 37C, 5% CO2. Then, 4 μL/well of ATPLite (PerkinElmer) was dispensed, incubated for 15 min at room temperature, and luminescence signal was measured using a Viewlux plate reader (PerkinElmer). Data was normalized with wells containing cells as 100%, and wells containing media only as 0% viability.

### Drug repurposing screen and data analysis

The NCATS pharmaceutical collection (NPC) was assembled internally, and contains 2,678 compounds, which include drugs approved by US FDA and foreign health agencies in European Union, United Kingdom, Japan, Canada, and Australia, as well as some clinical trialed experimental drugs [13]. The compounds were dissolved in 10 mM DMSO as stock solutions, and titrated at 1:5 for primary screens with 4 concentrations, and at 1:3 for follow up assays with 11 concentrations. The SARS-S PP entry assay in Vero E6 cells, and MERS-S PP entry assay in Huh7 cells, were used to screen the NPC library in parallel. Concurrently, counter screens for cytotoxicity of compounds in Vero E6 and Huh7 were also screened against the NPC library.

A customized software developed in house at NCATS [33] was used for analyzing the primary screen data. Half-maximal efficacious concentration (EC_50_) and half-maximal cytotoxicity concentration (CC_50_) of compounds were calculated using Prism software (GraphPad Software, San Diego, CA). Results in bar plots were expressed as mean of triplicates ± standard error of the mean (SEM).

### Luciferase immunofluorescence and high-content imaging

Cells were seeded at 15,000 cells in 100 μL/well media in 96-well assay plates, and incubated at 37□°C, 5% CO_2_ overnight (~16 h). Supernatant was removed, and 50 μL/well of PP was added. Plates were spin-inoculated at 1500 rpm (453 xg) for 45 min, incubated for 2 h at 37◻°C, 5% CO_2_, then 50 μL/well of growth media was added. The plates were incubated for 48◻h at 37◻°C, 5% CO_2_. Media was aspirated, and cells were washed once with 1X PBS (ThermoFisher). Cells were then fixed in 4% PFA (EMS) in PBS containing 0.1% BSA (ThermoFisher) for 30 min at room temperature. Plates were washed three times with 1X PBS, then blocked and permeabilized with 0.1% Triton-X 100 (ThermoFisher) in Cell Staining Buffer (Biolegend) for 30 min. Permeabilization/blocking solution was removed, 1:1000 primary mouse-anti-luciferase antibody (Santa Cruz) was added, and incubated overnight at 4 °C. Primary antibody was aspirated and cells were washed three times with 1X PBS. 1:1000 secondary antibody goat-anti-mouse-AlexaFluor 647 (ThermoFisher) was added for 1 h in Cell Staining Buffer. Cells were washed three times, and stained with 1:5000 Hoechst 33342 (ThermoFisher) and 1:10000 HCS Cell Mask Green (ThermoFisher) for 30 min, before three final 1X PBS washes. Plates were sealed and stored at 4 °C prior to imaging.

Plates were imaged on the IN Cell 2500 HS automated high-content imaging system. A 20x air objective was used to capture nine fields per well in each 96 well plate. Cells were imaged with the DAPI, Green, and FarRed channels. Images were uploaded to the Columbus Analyzer software for automated high-content analysis. Cells were first identified using the Hoechst 33342 nuclear stain in the DAPI channel. Cell bodies were identified using the HCS Cell Mask stain in the green channel using the initial population of Nuclei region of interests. Intensity of the FarRed channel indicating luciferase expression was measured, and a threshold was applied based on the background of the negative control. Average values, standard deviations, and data counts were generated using pivot tables in Microsoft Excel and data was plotted in Graphpad Prism.

### Negative stain and immunogold electron microscopy

All reagents were obtained from Electron Microscopy Sciences, unless otherwise specified. For negative staining without immunogold labeling, freshly glow-discharged, formvar and carbon coated, 300-mesh copper grids were inverted on 5 μl drops of sample on Parafilm for 1 min. Grids with adhered sample were transferred across two drops of syringe-filtered PBS, and then two drops of filtered distilled water before being placed on a drop of 1% aqueous uranyl acetate for 1 min, after which grids were blotted with filter paper, allowing a thin layer of uranyl acetate to dry on the grid.

SARS-S PP to be immunogold labeled were adhered to freshly glow discharged, formvar and carbon coated, 300-mesh gold grids, transferred across three drops of filtered PBS and then incubated on drops of filtered blocking solution containing 2% BSA (Sigma) in PBS for 10 min. Samples were covered during the incubation steps to prevent evaporation. Primary antibody to SARS-S (BEI), was diluted 1:20 in filtered blocking solution. After blocking, grids were blotted lightly with filter paper to remove excess solution before being transferred to primary antibody droplets and incubated for 30 minutes. Then, grids were transferred across two drops of blocking solution and incubated for 10 minutes. Secondary antibody (10 nm gold-conjugated Goat-α-Mouse IgG) was diluted 1:20 in filtered blocking solution. Grids were lightly blotted before transferred to droplets of secondary antibody, incubated for 30 min and then rinsed with 3 drops of PBS. Prior to negative stain, grids were transferred across three drops of distilled water to remove PBS as described above. Grids were observed using a ThermoFisher Tecnai T20 transmission electron microscope operated at 200 kV, and images were acquired using an AMT NanoSprint1200 CMOS detector (Advanced Microscopy Techniques).

### SARS-CoV-2 cytopathic effect (CPE) assay

SARS-CoV-2 CPE assay was conducted at Southern Research Institute (Birmingham, AL). Briefly, compounds were titrated in DMSO and acoustically dispensed into 384-well assay plates at 60 nL/well. Cell culture media (MEM, 1% Pen/Strep/GlutaMax, 1% HEPES, 2% HI FBS) was dispensed at 5 μL/well into assay plates and incubated at room temperature. Vero E6 (selected for high ACE2 expression) was inoculated with SARS CoV-2 (USA_WA1/2020) at 0.002 M.O.I. in media and quickly dispensed into assay plates as 4000 cells in 25 μL/well. Assay plates were incubated for 72 h at 37◻°C, 5% CO_2_, 90% humidity. Then, 30 μL/well of CellTiter-Glo (Promega) was dispensed, incubated for 10 min at room temperature, and luminescence signal was read on an EnVision plate reader (PerkinElmer). An ATP content cytotoxicity assay was conducted with the same protocol as CPE assay, without the addition of SARS-CoV-2 virus.

## Ethics Statement

The authors declared no potential conflicts of interest with respect to the research, authorship, and/or publication of this article.

## Author Contributions

MX, MP, KG, WZhu, MS and JP performed experiments; CZC and WZheng made the conceptualization and data curation; MS and GH analyzed data; CZC and WZheng wrote the manuscript, and all authors edited the manuscript.

## Acknowledgement

This work is funded by the Intramural Research Program of the National Center for Advancing Translational Sciences, and National Institute of Child Health and Human Development, National Institutes of Health. Work in the author’s laboratory (GW) is supported by the National Institutes of Health (research grant R01AI35270).

